# Sigflow: an automated and comprehensive pipeline for cancer genome mutational signature analysis

**DOI:** 10.1101/2020.08.12.247528

**Authors:** Shixiang Wang, Ziyu Tao, Tao Wu, Xue-Song Liu

**Affiliations:** School of Life Science and Technology, ShanghaiTech University, Shanghai 201203, China; Shanghai Institute of Biochemistry and Cell Biology, Chinese Academy of Sciences, Shanghai, China; University of Chinese Academy of Sciences, Beijing, China

## Abstract

**Summary:** Mutational signatures are recurring DNA alteration patterns caused by distinct mutational events during the evolution of cancer. In recent years, several bioinformatics tools are available for mutational signature analysis. However, most of them focus on specific type of mutation, or have limited scope of application. A pipeline tool for comprehensive mutational signature analysis is still lacking. Here we present Sigflow pipeline, which provides an one-stop solution for *de novo* signature extraction, reference signature fitting, signature stability analysis, sample clustering based on signature exposure in different types of genome DNA alterations including single base substitution (SBS), doublet base substitution (DBS), small insertion and deletion (INDEL), and copy number alteration. A Docker image is constructed to solve the complex and time-consuming installation issues, and this enables reproducible research by version control of all dependent tools along with their environments. The Sigflow pipeline can be applied to both human and mouse genomes.

**Availability and implementation:** Sigflow is an open source software under academic free license (AFL) v3.0 and it is freely available at https://github.com/ShixiangWang/sigminer.workflow or https://hub.docker.com/r/shixiangwang/sigflow.

## 1 Introduction

Mutational signatures reflect the accumulated effects of both exogenous and endogenous mutational processes acting on cancer cells. These specific patterns of mutational processes has been initially identified by Alexandrov and colleagues with nonnegative matrix factorization (NMF) based matrix decomposition algorithm in 2013 (Alexandrov, et al., 2013). Different other types of algorithms such as Bayesian NMF, Expectation–maximization have also been built for *do novo* mutational signature extraction (Baez-Ortega and Gori, 2019). The application of mutational signature analysis to ever-growing amount of sequencing data leads to the formation of COSMIC signature database (Alexandrov, et al., 2020).

Mutational signature analysis has been becoming a routine procedure after somatic variant calling in cancer genome study. This signature analysis can not only reveal the underlying mutational processes information, but also provide biomarkers for cancer precision stratification and clinical response prediction (Davies, et al., 2017; Ma, et al., 2018; Wang, et al., 2018). However, current available mutational signature analysis tools either provide limited analysis features, or only focus on specific type of genome alterations, such as single base substitutions (Baez-Ortega and Gori, 2019; Fischer, et al., 2013; Gehring, et al., 2015; Kim, et al., 2016; Mayakonda, et al., 2018; Rosenthal, et al., 2016). In addition, the installation and application processes of currently available tools are complex and time-consuming.

Here we present an open source pipeline tool Sigflow to provide a one-stop solution for efficient and reliable *de novo* signature extraction, reference signature fitting, signature exposure stability analysis, sample clustering based on signature exposure, etc. Sample level and signature level results are properly visualized. The SBS, DBS, INDEL signatures and the recently copy number signature analysis developed by our group are supported (Alexandrov, et al., 2020; Wang, et al., 2020). To solve the complex and time-consuming installation issues accompanying with current bioinformatics tools, a Docker image of Sigflow is constructed, and this enables good scalability for addition of other analysis features or other types of signatures in the future.

## 2 Tool description

Sigflow uses a command line based interface and allows the user to efficiently and automatically perform all the four workflows described below. The Sigflow (Fig. 1) begins with importing somatic variant data in MAF (recommended), VCF or CSV/EXCEL format, and then parses the user input to select the workflow to run. Subsequently, a sample by mutation catalogue matrix is generated. Finally, a user specified workflow is performed to extract and analyze mutational signatures. Important immediate and final results are saved to disk for general use. In addition, parallel computation is enabled, and this significantly reduces computation time.

**Fig. 1.**
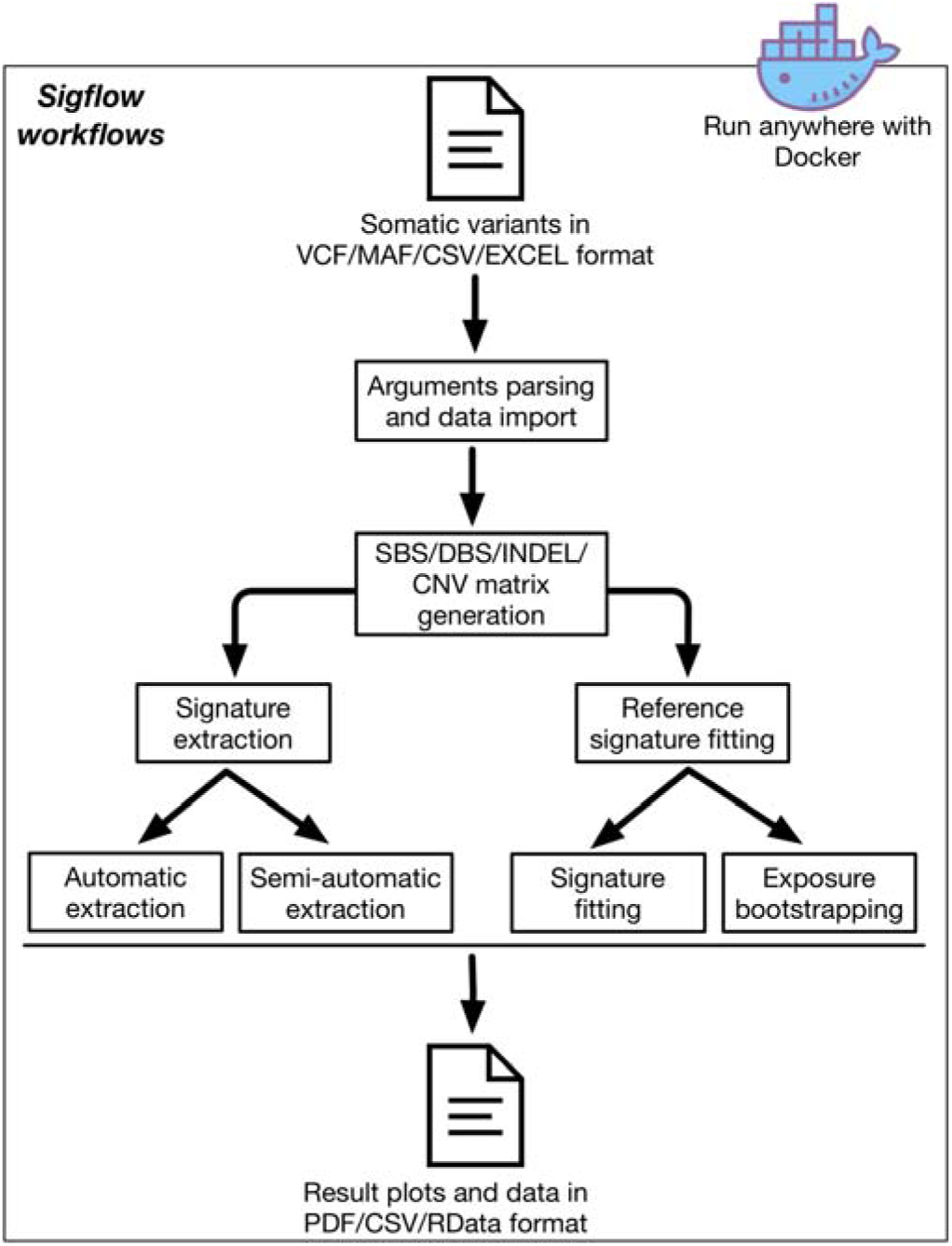
Overview of Sigflow pipeline.

### 2.1 Automatic *de novo* signature extraction

Two approaches are available in Sigflow for automatic *de novo* signature extraction. In the first approach, a Bayesian variant of NMF algorithm is applied to enable optimal inferences for the number of signatures through the automatic relevance determination technique (Kim, et al., 2016; Tan and Fevotte, 2013). This procedure starts from 30 signatures and reduces to a proper signature number which delivers highly interpretable and sparse representations for both signature profiles and exposures at a balance between data fitting and model complexity. The whole procedure can be run at a specified number of times and the most stable solution is selected as the final output. For example in total 10 runs, 5, 3 and 2 out of them support signature number 3, 4 and 5 respectively, then the best solution for signature number is 3. In the second approach, Sigflow directly calls SigProfiler, which is the original software for *de novo* mutational signature extraction (Bergstrom, et al., 2019). The SigProfiler results are collected and transformed into the same format as in the first approach. After extracting signatures, the data of signature profile and absolute/relative exposure are generated, samples are clustered by relative signature exposures, and cosine similarity analysis is performed to match the extracted signatures to the COSMIC reference signatures.

### 2.2 Semi-automatic *de novo* signature extraction

The Sigflow uses 2-step strategy in semi-automatic signature extraction. In the first step, it runs NMF at a specified number of times for signature number range from 2 to a proper max value (30 for SBS and copy number signature, 15 for DBS signature and 20 for INDEL signature; this value can be modified by the user), then outputs some common measures (e.g. cophenetic correlation coefficient, silhouette and residual sum of squares) for each signature number to help the user to determine the number of signature to extract (Gaujoux and Seoighe, 2010). The key point is to select a signature number which results in high reproducible mutational signatures and low overall reconstruction error (Alexandrov, et al., 2013). In the second step, Sigflow runs NMF at a specified number of times for the signature number from user input. Typically, 30 to 50 NMF runs can obtain a robust result (Gaujoux and Seoighe, 2010). This workflow has similar output files as the automatic signature extraction workflow.

### 2.3 Reference signature fitting

When the sample size is small (typically n<50), the d*e novo* workflows described above cannot properly decompose mutational signatures and their exposures. To extract signatures from single sample, an algorithm was designed to find a linear combination of the predefined signatures (such as COSMIC signatures) that best reconstructs the sample’s mutational profile. Here Sigflow uses quadratic programming algorithm for reference signature fitting, and this algorithm is originally implemented in SignatureEstimation package and is fast and reliable (Huang, et al., 2018). This workflow is computationally efficient (typically finished in several minutes for 100 samples) and is recommended for input data with small sample size.

### 2.4 Signature exposure stability analysis

The results from different signature analysis methods are not always consistent. Hence, one needs to be able to not only decompose a patient’s mutational profile into signatures but also establish the accuracy of such signature decomposition (Huang, et al., 2018). Bootstrapping analysis is performed to quantify the confidences in the estimated exposure of each mutational signature. By randomly re-sampling original mutational catalogs for each tumor sample at a specified number of times, this workflow generates estimated bootstrap confidence intervals for each signature exposure and computes an empirical probability (p value) that a relative signature exposure is above a specific threshold. Signature exposure instability is also measured as the root mean squared error (RMSE) of the exposure differences between bootstrap estimates and the optimal solutions in the original data to test how much the bootstrap exposures vary from original exposures. The outputs of this analysis include bootstrap exposures, reconstruction errors and p values under different exposure cutoffs.

## 3. Implementation

The Sigflow pipeline tool has been developed with R 4.0 following a clean, modular and robust design in concordance with best practice coding standards. Instructions on how to install and run the Sigflow are presented in the public GitHub repository (https://github.com/ShixiangWang/sigminer.workflow) as well as Supplementary file. A detailed manual, which describes the workflows and operating parameters, is also provided in the GitHub README page and Supplementary file. The Sigflow is highly customizable with numerous parameter settings and is well supported for different input file formats, and all options are explained in the integrated help section. It has been designed to run as a command line based program with a user-friendly interface, which allows non-expert users to become quickly familiarized. The Sigflow allows keeping R related data files, which can be easily loaded into R for flexible and interactive analysis and visualization by Sigminer and other R packages.

To enable quick and reproducible research, we built a version-controlled Docker image for Sigflow to avoid the complex and time-consuming dependency issues in bioinformatics tool installation. Due to the flexibility of container technology, Sigflow can be easily deployed, managed and deleted on any operating system, thus it is convenient to be integrated with other cancer genome analysis platforms.

## 4 Conclusion

In the recent years we have witnessed an increased number of tools and studies that explore and utilize mutational signatures in different aspects, including mutational etiologies exploration, biomarker discovery and cancer evolution. For better data integration and explanation, and higher computational efficiency, it is important to build robust, efficient and user-friendly tool that eventually allow a wide range of users to perform mutational signature analysis. The Sigflow is a novel pipeline tool that provides comprehensive mutational signature analysis workflows, supports easy and quick tool deployment, and reproducible research.

## Acknowledgements

We thank ShanghaiTech University High Performance Computing Public Service Platform for computing services.

## Funding

This work was supported in part by The National Natural Science Foundation of China (31771373), and startup funding from ShanghaiTech University.

## Conflict of Interest

none declared.

## Notes

### Competing Interest Statement

The authors have declared no competing interest.

## Reference

Alexandrov, L.B., et al. The repertoire of mutational signatures in human cancer. Nature 2020;578(7793):94–101.

Alexandrov, L.B., et al. Deciphering signatures of mutational processes operative in human cancer. Cell Rep 2013;3(1):246–259.

Baez-Ortega, A. and Gori, K. Computational approaches for discovery of mutational signatures in cancer. Brief Bioinform 2019;20(1):77–88.

Bergstrom, E.N., et al. SigProfilerMatrixGenerator: a tool for visualizing and exploring patterns of small mutational events. BMC Genomics 2019;20(1):685.

Davies, H., et al. HRDetect is a predictor of BRCA1 and BRCA2 deficiency based on mutational signatures. Nat Med 2017;23(4):517–525.

Fischer, A., et al. EMu: probabilistic inference of mutational processes and their localization in the cancer genome. Genome Biol 2013;14(4):R39.

Gaujoux, R. and Seoighe, C. A flexible R package for nonnegative matrix factorization. BMC Bioinformatics 2010;11:367.

Gehring, J.S., et al. SomaticSignatures: inferring mutational signatures from single-nucleotide variants. Bioinformatics 2015;31(22):3673–3675.

Huang, X., Wojtowicz, D. and Przytycka, T.M. Detecting presence of mutational signatures in cancer with confidence. Bioinformatics 2018;34(2):330–337.

Kim, J., et al. Somatic ERCC2 mutations are associated with a distinct genomic signature in urothelial tumors. Nat Genet 2016;48(6):600–606.

Ma, J., et al. The therapeutic significance of mutational signatures from DNA repair deficiency in cancer. Nat Commun 2018;9(1):3292.

Mayakonda, A., et al. Maftools: efficient and comprehensive analysis of somatic variants in cancer. Genome Res 2018;28(11):1747–1756.

Rosenthal, R., et al. DeconstructSigs: delineating mutational processes in single tumors distinguishes DNA repair deficiencies and patterns of carcinoma evolution. Genome Biol 2016;17:31.

Tan, V.Y. and Fevotte, C. Automatic relevance determination in nonnegative matrix factorization with the beta-divergence. IEEE Trans Pattern Anal Mach Intell 2013;35(7):1592–1605.

Wang, S., et al. APOBEC3B and APOBEC mutational signature as potential predictive markers for immunotherapy response in non-small cell lung cancer. Oncogene 2018;37(29):3924–3936.

Wang, S., et al. Copy number signature analyses in prostate cancer reveal distinct etiologies and clinical outcomes. medRxiv 2020.

