## Supplemental Figure and note for "Sigflow: an automated and comprehensive pipeline for cancer genome mutational signature analysis"

### Supplementary materials for Sigflow

#### Supplementary figures

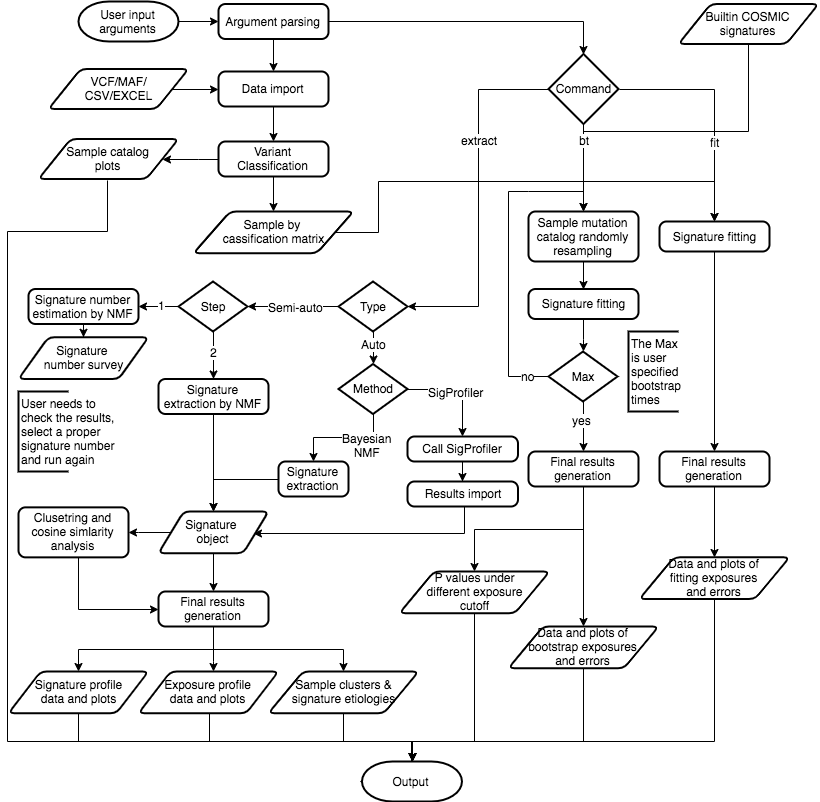

Figure S1. Detailed workflow of Sigflow.

#### Supplementary software documentation

##### Installation

Sigflow can be installed in any operating system (Windows/MacOS/Linux). The user can choose one of the following two ways to install Sigflow.

- Step-by-step installation.
- Install Sigflow Docker image.

###### Step-by-step installation

This process takes about 1 hour to finish (depends on your internet speed). This is recommended for R users.

1. Install R from [CRAN](https://cran.r-project.org/) (R >3.6 is required).
2. Install R packages by running the following commands in your R console.

- install.packages("docopt")
  install.packages("BiocManager")
  BiocManager::install("sigminer", dependencies = TRUE)
  ## Install specific version of sigminer
  ## e.g. sigminer v1.0.10
  # install.packages("remotes")
  # remotes::install_github("ShixiangWang/sigminer@v1.0.10")

3. (Optional) Install required reference genome R packages. The hg19 genome build is installed along with package sigminer. To install hg38 or mm10 genome build, run the following R commands.

- # For hg38
  BiocManager::install("BSgenome.Hsapiens.UCSC.hg38")
  # For mm10
  BiocManager::install("BSgenome.Mmusculus.UCSC.mm10")

4. Clone Sigflow GitHub repository.

- $ git clone https://github.com/ShixiangWang/sigminer.workflow

5. (Optional) Link the R script as a executable file.

- $ cd sigminer.workflow
  $ ln -s $PWD/sigflow.R /usr/bin/sigflow # You can choose another place instead of /usr/bin/sigflow

If you have the the step 5, you can restart your terminal and check the Siglow command by running siglow -h, otherwise you can directly run this script by ./sigflow.R -h.

###### Install by Docker

This process takes about 10 minutes to finish (depends on your internet speed). This is recommended for non-R users or users have no programming experience.

1. Follow Docker [official documentation](https://docs.docker.com/) to install Docker.
2. Install latest stable version of Sigflow docker image by running the following command in your terminal.

- $ sudo docker pull shixiangwang/sigflow:1.0
- Here $ is the prompt.

Now you can run Siglow using the following command.

$ docker run shixiangwang/sigflow

##### Usage

The **Installation** section has described how to install Sigflow and call it.

Sigflow now provides 4 workflows which implemented in 3 commands:

- extract
- fit
- bt

The details of each command and corresponding flags are documented as below.

```
=================================================================
sigflow: Streamline Analysis Workflows for Mutational Signatures.

Author: Shixiang Wang
Author: Xue-Song Liu
Copyright: AFL@2020 [https://opensource.org/licenses/AFL-3.0]

Desc:
  There are several subcommands.
  ==
  extract - extract signatures by either automated or manual way.
            Of note, when you use manual way, you need to run 2 times,
            firstly you should set --manual to get signature estimation results,
            and secondly you should set --manual --number N to get N signatures.
  ==
  fit     - fit signatures in >=1 samples based on COSMIC reference signatures.
  ==
  bt      - run bootstrap signature fitting analysis in >=1 samples.

Usage:
  sigflow extract --input=<file> [--output=<outdir>] [--mode=<class>] [--manual --number <sigs>] [--max <max>] [--genome=<genome>] [--nrun=<runs>] [--cores=<cores>] [--sigprofiler] [--hyper] [--verbose]
  sigflow fit --input=<file> [--output=<outdir>] [--mode=<class>] [--genome=<genome>] [--verbose]
  sigflow bt  --input=<file> [--output=<outdir>] [--mode=<class>] [--genome=<genome>] [--nrun=<runs>] [--verbose]
  sigflow (-h | --help)
  sigflow --version

Options:
  -h --help     Show help message.
  --version     Show version.
  -i <file>, --input <file>       input file/directory path.
  -o <outdir>, --output <outdir>  output directory path [default: ./sigflow_result/].
  -m <class>, --mode <class>      extract/fit mode, can be one of SBS, DBS, ID, MAF (for three types), CN (not supported in fit subcommand) [default: SBS].
  --manual                        enable manual extraction, set -N=0 for outputing signature estimation firstly.
  -N <sigs>, --number <sigs>      extract specified number of signatures [default: 0].
  --max <max>                     maximum signature number, default is auto-configured, should >2 [default: -1].
  -g <genome>, --genome <genome>  genome build, can be hg19, hg38 or mm10, [default: hg19].
  -r <runs>, --nrun <runs>        run times of NMF (extract) or bootstrapping (bt) to get results [default: 30].
  -T <cores>, --cores <cores>     cores to run the program, large dataset will benefit from it [default: 1].
  --hyper                         enable hyper mutation handling in COSMIC signatures (not used by SigProfiler approach).
  --sigprofiler                   enable auto-extraction by SigProfiler software.
  -v, --verbose                   print verbose message.

=================================================================
```

Sigflow supports input data in VCF/MAF/CSV/EXCEL format. The file format is auto-detected by Sigflow.

For SBS/DBS/INDEL data in CSV (including TSV) or EXCEL format, the following columns typically described in MAF format are necessary:

- 'Hugo_Symbol': gene symbol
- 'Chromosome': chromosome name, e.g. "chr1"
- 'Start_Position': start positionof the variant (1-based)
- 'End_Position': end position of the variant (1-based)
- 'Reference_Allele': reference allele of the variant, e.g. "C"
- 'Tumor*Seq*Allele2': tumor sequence allele, e.g. "T"
- 'Variant*Classification': variant classification, e.g. "Missense*Mutation"
- 'Variant_Type': variant type, e.g. "SNP"
- 'Tumor*Sample*Barcode': sample identifier

For copy number segment data in in CSV (including TSV) or EXCEL format, the following columns are necessary:

- 'Chromosome': chromosome name, e.g. "chr1"
- 'Start.bp': start breakpoint position of segment
- 'End.bp': end breakpoint position of segment
- 'modal_cn': integer copy number value
- 'sample': sample identifier

##### Suggestion and bug report

Any suggestion, feature request or bug report is welcome at [GitHub repository issue](https://github.com/ShixiangWang/sigminer.workflow/issues).

##### Examples

Example datasets along with many example code are available in clone repository above (you can read it online at [here](https://github.com/ShixiangWang/sigminer.workflow/tree/master/test)).

The following parts give an example for each command.

Result directory of any command has the following structure.

- Files with extension.RData and .rds are R related files to reproduce the results, and can be imported into R for further analysis and visualization.
- Files with extension.pdf are common visualization results used for communication.
- Files with extension .csv are formated data tables used for inspection, communication or further analysis.

###### extract command

$ # Assume you have done the clone step
$ # git clone https://github.com/ShixiangWang/sigminer.workflow
$ cd sigminer.workflow/test
$ sigflow extract -i tcga_laml.maf.gz -o test_results/test_maf -m MAF -r 10 -T 4 --max 10

This will auto-extract SBS/DBS/INDEL signatures from data toga_laml.maf.gz by 10 Bayesian NMF runs with 4 computer cores (4 threads) from signature number ranges from 1 to 10, output results to directory test_results/test_maf.

###### fit command

$ # Assume you have done the clone step
$ # git clone https://github.com/ShixiangWang/sigminer.workflow
$ cd sigminer.workflow/test
$ sigflow fit -i tcga_laml.maf.gz -o test_results/test_fitting -m MAF

This will auto-fit input data tcga_laml.maf.gz to COSMIC SBS/DBS/INDEL signatures. Signature exposure data tables and plots are outputed.

###### bt command

$ # Assume you have done the clone step
$ # git clone https://github.com/ShixiangWang/sigminer.workflow
$ cd sigminer.workflow/test
$ sigflow bt -i tcga_laml.maf.gz -o test_results/test_bt -m SBS -r 5

This will auto-fit the random resample of input mutation profile to COSMIC SBS/DBS/INDEL signatures for specified times (here is 5). Data tables and plots of bootstrap signature exposures, errors and p values under different exposure cutoff are outputed.

NOTE, in practice, set -r to a value >=100 is recommended.

###### How to use Docker to run Sigflow

If you use Docker to run Sigflow, you cannot directly call sigflow command. Instead, you should use sudo docker run --rm -v /your_local_path:/docker_path shixiangwang/sigflow to start a Docker container.

For example, if you want to accomplish the same task shown in extract command above, you need to run:

$ sudo docker run --rm -v /your_local_path:/docker_path shixiangwang/sigflow extract -i /docker_path/tcga_laml.maf.gz -o /docker_path/test_maf -m MAF -r 10 -T 4 --max 10

Here,

- --rm will delete this container when this task is finished.
- -v is used for mounting your local directory /your_local_path as /docker_path in Docker image. **This is important**. You need to use the Docker container path in Sigflow arguments. So there must be a file called /your_local_path/tcga_laml.maf.gz exists in your computer, it will be treated as /docker_path/test_maf in the container.
